# Plastic cell morphology changes during dispersal

**DOI:** 10.1101/2021.06.13.448130

**Authors:** Anthony D. Junker, Staffan Jacob, Hervé Philippe, Delphine Legrand, Chad G. Pearson

## Abstract

Dispersal is the movement of organisms from one habitat to another that potentially results in gene flow. It is often found to be plastic, allowing organisms to adjust dispersal movements depending on environmental conditions. A fundamental aim in ecology is to understand the determinants underlying dispersal and its plasticity. We utilized 22 strains of the ciliate *Tetrahymena thermophila* to determine if different phenotypic dispersal strategies co-exist within a species and which mechanisms underlie this variability. We quantified the cell morphologies impacting cell motility and dispersal. Distinct differences in innate cellular morphology and dispersal rates were detected, but no universally utilized combinations of morphological parameters correlate with dispersal. Rather, multiple distinct and plastic morphological changes impact cilia-dependent motility during dispersal, especially in proficient dispersing strains facing challenging environmental conditions. Combining ecology and cell biology experiments, we show that dispersal can be promoted through a panel of plastic motility-associated changes to cell morphology and motile cilia.

**Graphical abstract:** 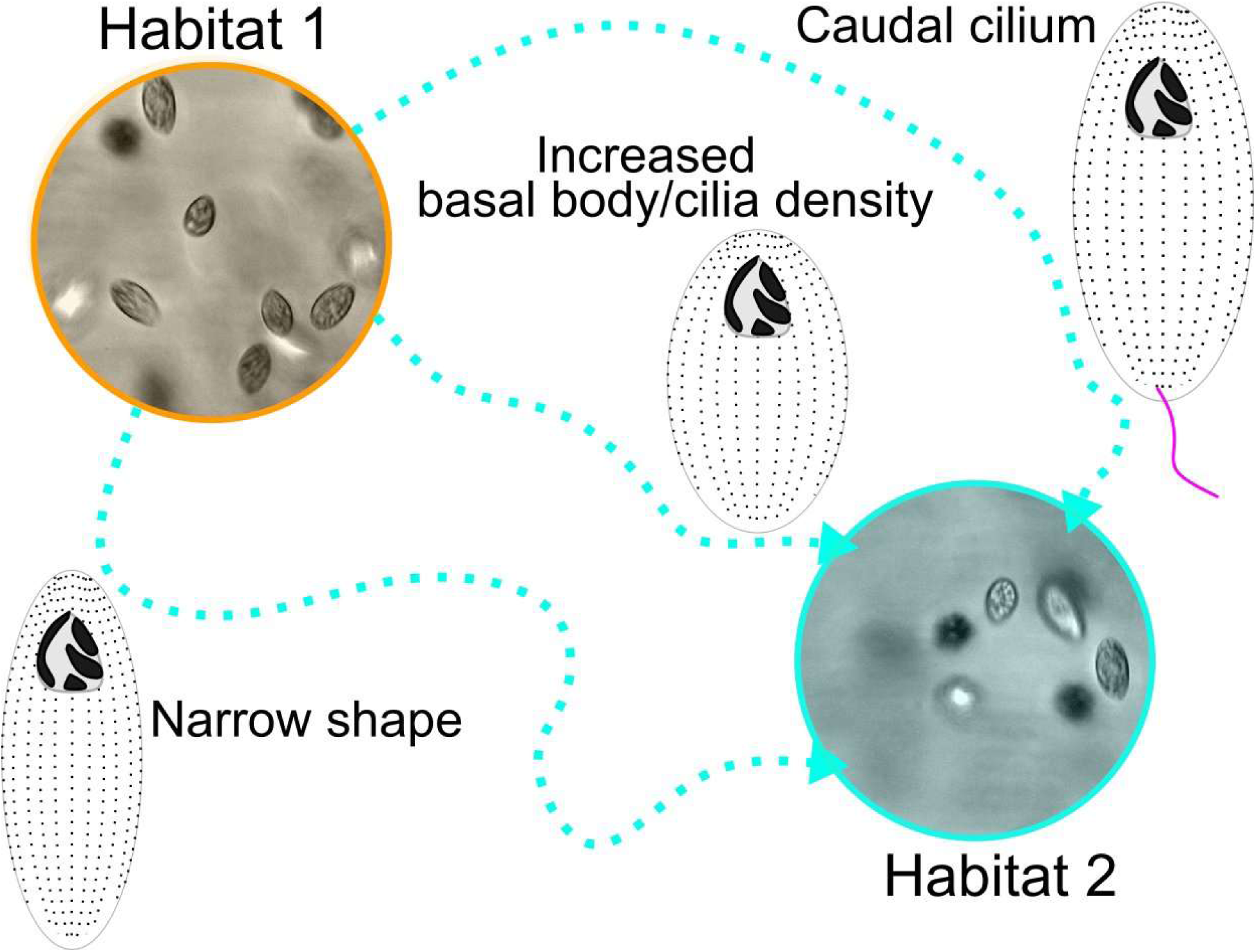

**Highlights:**

- *Tetrahymena thermophila* exhibits intra-specific diversity in morphology and dispersal.
- Cell motility behavior during dispersal changes with cilia length and cell shape.
- Cells from proficient dispersing strains transiently change basal body and cilia position.
- Starvation-induced dispersal triggers increased basal body and cilia density and caudal cilium formation in rapid-swimming cells.

## Introduction

Organisms move within or between habitats in the search for food or partners, or to avoid unsuitable local conditions^1^. Routine movements occur mostly within habitats, while a movement between habitat patches influencing gene flow from one population to another is defined as dispersal^2, 3^. During dispersal, an organism leaves a local habitat patch (emigration), moves through a matrix separating habitats (transit), and finally colonizes or settles in a new habitat (immigration). Dispersal movements often depend upon both the environmental conditions encountered at each of its three phases (*e.g.*, density- or resource-dependent dispersal), and upon the phenotype of the individuals^4^. Dispersing individuals can be phenotypically different from residents for a suite of correlated traits. This correlation between the dispersal rate and phenotypic traits defines a dispersal syndrome^4, 5^. Traits involved in dispersal syndromes can be diverse, linked to morphology, behavior, physiology and/or life-history^5^. Moreover, dispersal syndromes have been described in a variety of taxa including plants, fish, rodents, and arthropods^4, 6–10^.

The acquisition of a repertoire of dispersal strategies may enhance a species’ ability to cope with the spatio-temporal heterogeneity of environmental conditions^11^. However, in most organisms, neither the extent of intra-specific variability in dispersal syndromes^6, 7^, nor the underlying mechanisms behind this diversity (genetic variation or phenotypic plasticity) are deeply understood^12^. Depending on these underlying mechanisms, the direction and speed of response to environmental changes diverges. Plastic strategies can facilitate the rapid fine-tuning of dispersal-related traits of a single genotype to environmental conditions but correspond with the costs of expressing such phenotypes^13^. Plasticity (including that which is linked to dispersal) can also limit local adaptation to new environmental conditions in cases where it buffers the phenotypic variation of genetically-diversified populations because it hinders the diversity of existing genotypes to the effects of natural selection^14^. Conversely, genetically-driven dispersal strategies may result from a long adaptive history of the species to its environmental landscape without paying the cost of environment-dependent strategies. But genetic diversity might not be sufficient for a population to face rapid and drastic environmental shifts. Evaluating the extent of phenotypic variation in dispersal syndromes that rely on both genetic and plastic mechanisms is important to capture the consequences of dispersal strategies on ecological and evolutionary dynamics, especially in the face of current global change^1, 15^.

Understanding the morphological and behavioral changes during dispersal requires their quantitative observation at corresponding spatial and temporal scales. Microbes are tractable for dispersal behavior and morphology studies because of their small sizes, short generation times, and morphologies and motilities that can now be readily observed^16^. The ciliate, *Tetrahymena thermophila*, lives in freshwater ecosystems and undergoes environmentally stimulated morphological and motility dynamics^17–19^. *T. thermophila* isolates exhibit context-dependent dispersal behaviors in response to crowding, nutrient availability, presence of kin, and temperature^20–25^. *T. thermophila* cells swim by the coordinated beating of hundreds of motile cilia that cover the approximately 20 x 40 µm rugby ball shaped cell. Cell shape is dictated, in part, by the aspect ratio of length relative to width. Ciliary beating produces hydrodynamic flow along the cell shape and is essential for both directed cell swimming and tortuous navigation through the cell’s environment^26^. Cilia are nucleated and positioned by basal bodies (BBs). Importantly, changes to cilia length and their positioning directly influence both ciliary motility and cell morphology^27–31^. In the case of cilia length, intraflagellar transport (IFT) defects cause cilia to shorten, reducing cell motility and eventually paralyzing cells^28, 32, 33^. In the case of BB and cilia position, loss of BB stability factors reduces BB and cilia number in ciliary rows and also slows cellular motility^34–36^. Furthermore, disrupting BB anchoring structures causes BBs to be mispositioned, which alters cell morphology and reduces the speed and linearity of cell swimming^27, 37, 38^. In contrast, starvation of *T. thermophila* results in highly motile cells with behaviors (fast and directed swimming) suggested to be associated with dispersal. Upon starvation, cells halt the cell division cycle, narrow in cell shape, increase ciliary density, and increase their ciliary beating rate^24, 39, 40^. These changes increase the rate of cell motility. Thus, morphology changes to *T. thermophila* cells are likely associated with dispersal behaviors.

In *T. thermophila*, dispersing cells generally take the form of smaller, more elongated, and fast-swimming cells as a result of plastic mechanisms^41^, although the morphologies of dispersers can vary among genotypes (e.g., large, round cells^21^, see also^24^). This variability in gross morphology associated with cell shape (aspect ratio) and ciliary spacing influences the dynamics of spatially structured cell population but has not been studied in detail^21, 24, 41–43^. We hypothesize that the described dispersal syndromes and their variability result from mechanistic changes at BBs and cilia that impact cell morphology and motility. Here, we define the link between fine-scale cellular processes and dispersal using both innate (genetic and stable non-genetic factors) and plastic (non-genetic factors sensitive to the environment) variation in behavior and cell morphology between strains of *T. thermophila*.

We used an experimental approach to relate the dispersal rates of 22 clonal strains of *T. thermophila* with both innate and plastic cell morphologies impacting cell motility. Given the diversity of documented dispersal responses of this species when exposed to either standard or stressful conditions, we hypothesize that several morphological, especially cilia-dependent, and behavioral dispersal strategies co-exist in *T. thermophila*^13, 22–24^. We first used two-patch systems commonly used to study dispersal under non-challenging environmental conditions to decipher the basics of dispersal phenotypic determinants^21, 41–43^. We then exposed the most and least dispersive strains to starvation, a stressful environmental condition described as a driver of strong dispersal response, and measured the same phenotypic metrics as in standard conditions^24, 39^. By bridging the usually unrelated evolutionary ecology and cell biology fields, we reveal an unexpected repertoire of dispersal strategies that rely on both innate and plastic phenotypes, opening new perspectives in the extent of intra-specific dispersal syndrome variability.

## Results

### *Tetrahymena thermophila* isolates exhibit unique dispersal propensities

We first leveraged dispersal rate variation in 22 clonal *T. thermophila* strains under standard conditions (D1 to D22, Figure S1A). Using a system of two habitat patches connected by a corridor tube (Figure 1A), cells were seeded into one habitat patch and allowed to disperse to the second patch through the corridor over 4 hours (less than one asexual generation; time interval allows a representative estimate of a strain dispersal rate, V. Thuillier and N. Schtickzelle, unpublished). This routinely used setup allows for discrimination between *T. thermophila* ‘resident’ cells that remain in the original patch and ‘dispersing’ cells that move to the second patch^44^. Cell movement might lead to a homogenous distribution across the system within a few minutes after inoculation^45^. However, this does not occur, meaning that movements between patches clearly deviates from random cell diffusion and instead result from dispersal decisions^45^. The proportion of dispersing cells was used to define the dispersal rate. Most of the strains fell within one standard deviation of the mean dispersal fraction (Figure 1B). However, strains harboring both low and high dispersal propensities were observed. D4 is the lowest dispersing strain with 29% of the cells dispersed to the secondary chamber. Conversely, D5 is the highest dispersing strain with 88% of the cells dispersed to the secondary chamber. This amounts to a threefold range in dispersal rates across the 22 strains. Thus, *T. thermophila* show differing dispersal propensities, which may inform the rates at which strains move between habitat patches in nature.

**Figure 1.**
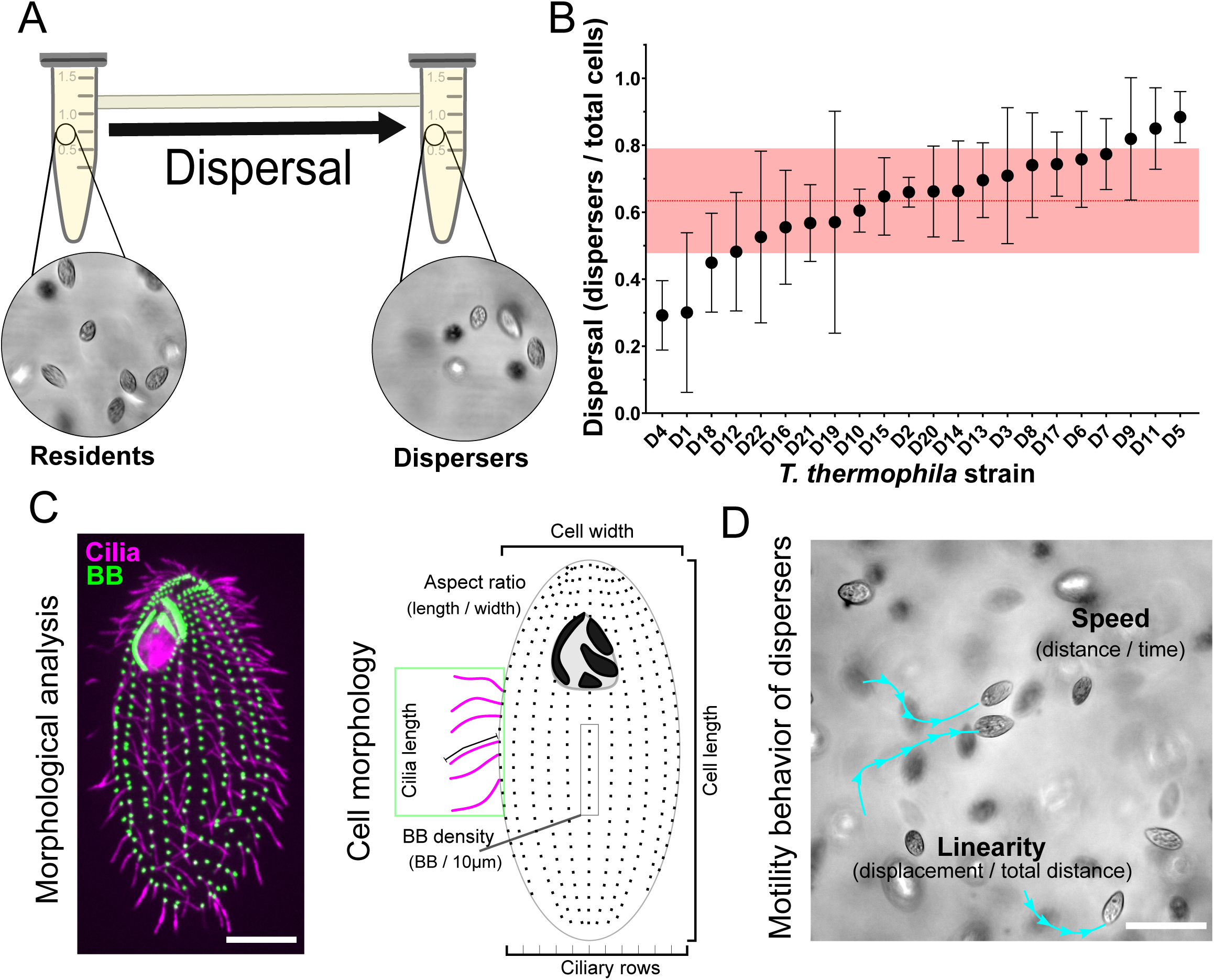
Differential dispersal propensities in *T. thermophila* strains. (A) Schematic of the two-patch dispersal system composed of 1.5 mL conical tubes connected by a narrow passageway. Cells in dispersal assays are seeded in the left patch and allowed to disperse to the right patch over 4 hr. (B) Dispersal propensity is different between the 22 *T. thermophila* isolates. Dispersal rate is measured as a ratio between number of dispersers and total number of cells in 5 separate biological replicates. Graph shows the mean and standard deviation in dispersal rate for each strain. Red line and transparent red band denote the inter-strain mean and inter-strain standard deviation for all strains. (C) Fluorescence image of *T. thermophila* cell with labeled BBs (α-TtCen1; green) and cilia (α-α-tubulin; magenta) (scale bar, 10 µm). Schematic of cell morphologies that were measured in the 22 *T. thermophila* isolates. (D) Schematic for *T. thermophila* cell behavior measurements (scale bar, 10 µm).

### Diverse innate morphologies and motilities of *Tetrahymena thermophila*

To determine if innate cell morphologies associated with cell motility could underlie the variability in dispersal propensity observed among strains, we quantified the distribution of cell and ciliary morphologies in the 22 strains under uniform growth conditions (Figure 1C, Figures S1 and S2) and compared them with the laboratory strain B2086. Mean ± standard deviation measurements from 19.7±1.7 cells were generated for each strain and compared to the cumulative inter-strain mean for each parameter. Each strain was then plotted relative to the inter-strain average to describe their cell morphology. The length of all 22 strains was 42.0±5.0 µm and the width was 20.0±2.9 µm. The aspect ratio (length/width) was 2.1±0.4. The BB, and therefore cilia density, was 7.1±0.4 BBs / 10 µm. The number of ciliary rows was 20.0±1.9 rows. The cilia length was 5.5±0.4 µm. Finally, the swim speed was 244±96 µm/sec. The laboratory strain B2086 closely resembled the inter-strain means but exhibited 97% less variability compared to the 22 strains (Figure S2B and S2C). Overall, the 22 isolates exhibited a broad spectrum of morphological variation from the inter-strain mean.

### Innate cell shape and cilia length inform swimming linearity in dispersers

Motility of organisms from one habitat to another is an essential component of dispersal, and the BB and cilia parameters that promote cilia-dependent cell propulsion is important for cell motility ^27, 30, 37, 46, 47^. First, we tried to identify innate patterns of trait associations across the 22 strains that could be linked to dispersal propensity based on the set of traits described in the previous section. Second, we added dispersers’ swimming behaviors (swim speed and linearity of cell having moved to the second patch, Figure 1D) to the analysis in the aim to find correlations between innate morphological traits and dispersers’ behavior. The morphological analysis of the three most and three least proficient dispersing strains are shown as examples (Figure 2A). Each measured parameter was ranked among the 22 strains and simple linear regression was used to compare the parameters (Figure 2B).

**Figure 2.**
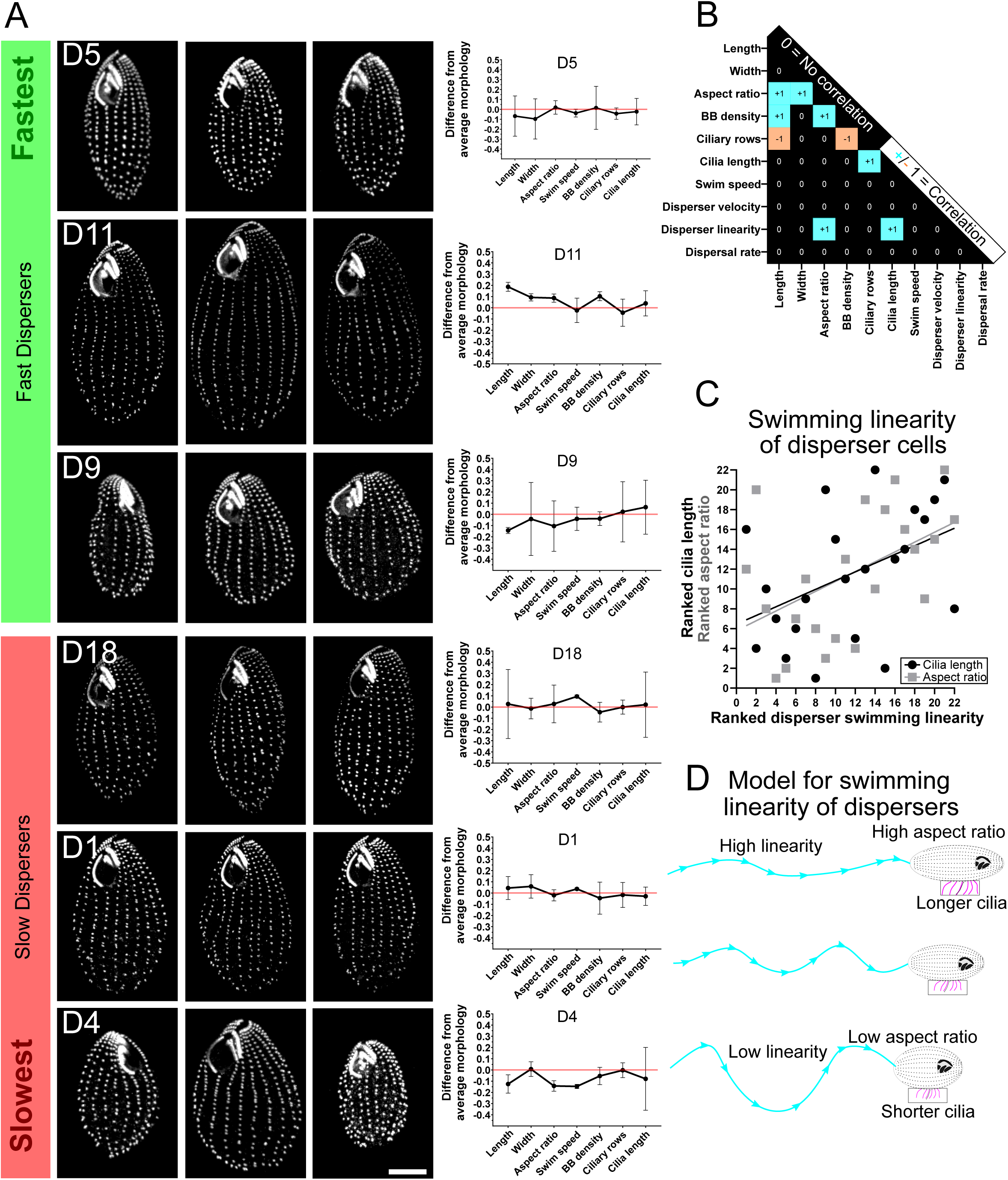
Increased linearity of swimming in high disperser cells correlates with narrow cell shape and longer cilia. (A) Fluorescence images of BBs (α-TtCen1; greyscale) and cell morphology from the three highest and three lowest dispersing strains among the 22 *T. thermophila* strains (scale bar, 10 µm). Graphs indicate the individual mean and standard deviation for each morphology parameter relative to the mean morphology for all strains (red line). (B) Plot of correlations between the ranked morphologies and the dispersal rate of the 22 *T. thermophila* strains. Correlation was calculated using linear regression between morphologies. Correlation is defined as F-test non-zero slopes of p=<0.05 denoted by a boxed blue “+1” (positive correlation) or orange “-1” (negative correlation) and no correlation is defined as F-test non-zero slopes of p=>0.05 denoted by a black boxed by “0” (see Methods). (C) Plot of the ranked innate aspect ratio (cell length / width) and innate cilia length relative to ranked swimming linearity (displacement/total distance) in disperser cells. Linear regression shows slopes are non-zero for aspect ratio (p=0.04) and cilia length (p=0.03). Higher ranked numbers (0-22) denote increased ranked linearity, increased ranked aspect ratio, and longer ranked cilia. (D) Model of *T. thermophila* cell morphology relative to swimming behavior during dispersal.

Innate cell aspect ratio significantly correlated with cell length and width (R^2^=0.47, p=<0.01 and R^2^=0.28, p=0.01, respectively), as expected. Tetrahymena lab strains exhibit a counting mechanism that maintains a constant number of BBs per cell^17, 48^. Changes to BB density and the number of ciliary rows was observed to impact cell length and width. Consistently, we showed in the 22 strains that BB density predicted cell length and the number of ciliary rows such that increased BBs density correlated with increased cell length and reduced number of total rows (Figure 2B). However, the number of ciliary rows did not correlate with cell width as previously observed with *T. thermophila* lab strains^17, 48^. This is because several strains exhibited unique morphologies when comparing ciliary row number and cell width. D9 for instance had many ciliary rows (20.2) relative to its narrow width (18.9 µm) and D11 a low frequency of ciliary rows (18.6) but was very wide (22.0 µm). This suggests that parameters beyond BB and ciliary placement control cell size and shape. In addition, cilia length increases in cells with more ciliary rows (Figure 2B).

The linearity of swimming in cells that have moved to the second patch (dispersers) separately correlated with innate cell aspect ratio and innate cilia length (Figure 2C; p<0.04). Dispersers that swim more linearly have longer cilia and/or have narrow cell shapes (high aspect ratio, Figure 2D). In contrast, dispersers that swim less linearly have shorter cilia and/or have ‘fatter’ cell shapes (low aspect ratio). Cilia length and cell shape do not independently correlate with each other (Figure 2B). Thus, strains with differing innate morphologies employ differing behavioral strategies during dispersal.

### Transient increase in BB density and cell shortening correspond with dispersal

In addition to innate factors as shown in the previous section, dispersal can be modulated by plastic mechanisms. We tested if dispersal propensity was impacted by intra-generational plasticity in cell morphology during the dispersal process by comparing resident and disperser morphologies. Such cell morphology differences after dispersal assays in isogenic strains can indeed be explained by cells plastically expressing new morphologies when confronted with dispersal (increase in global phenotypic variance). However, it can also be explained by morphological subpopulations in the starting population that sort into residents and dispersers during dispersal (equal global phenotypic variances), or by selective mortality of morphological subpopulations during dispersal (decrease in global phenotypic variance). We thus additionally compared trait distributions in starting populations to residents and dispersers to exclude these two last scenarios.

We detailed changes in cell morphologies between dispersers and residents 0, 180 and 360 min after dispersal induction in three strains. We leveraged the spectrum of low (D4), high (D5) and very high (B2086) dispersing strains to ask how plastic changes to morphology is linked to dispersal propensity (Figure 3 and S3). Clear plastic changes between residents and dispersers were not observed for most morphological parameters (Figure S3 J-K). However, consistent with the relationship between cell shape and BB spacing found above, we identified a relationship between cell length and BB density in disperser cells. While there was a general decrease in cell length at 180 and 360 min amongst all cells (Figure 3A), dispersers showed a more pronounced decrease compared to residents. Dispersers from D4, a low dispersing strain, showed a 7% and 5% decrease in cell length, respectively after 180 and 360 min compared to residents. D4 BB density remained unchanged at each time point (Figure 3B). Dispersers from D5, a highly dispersing strain, had a more pronounced decrease in cell length at 180 (13%) and 360 (14%) min compared to resident cells. B2086 disperser cell length was unchanged at 180 min but shortened at 360 min (4%). Consistent with the shorter cells, BB density in disperser cells increased at 180 (10% and 6% for D5 and B2086, respectively) and 360 (16% and 10%) min compared to residents. The distribution of morphologies in cell length and BB density of residents and dispersers extend outside the range of morphology displayed in the starting population (Figure 3A,B). The overlap of the start morphologies to the respective means in the dispersal systems are lowest for cell length in the D5 (6% overlap) and B2086 (8% overlap) t360 dispersers, as compared to the D4 (19% overlap) t360 dispersers (Figure S3G,H). The overlap of the start conditions to the respective means in simulations are lowest for BB density in the D5 (4% overlap) and B2086 (10% overlap) t360 residents, as compared to the D4 (21% overlap) t360 residents. Thus, the innate morphologies do not appreciably contribute to the mean morphology observed in resident and dispersal populations. Moreover, the standard deviation is increased in residents and dispersers in higher dispersing cells compared to their respective start conditions (D5 and B2086 cells (Figure S3E,F). This suggests morphological plasticity in residents and dispersers, rather than sorting of innate starting population morphologies or selective mortality, best explains the decreased in cell length and increased BB density in cells with higher dispersal. This highlights the importance of transient morphological changes to support motility during dispersal.

**Figure 3.**
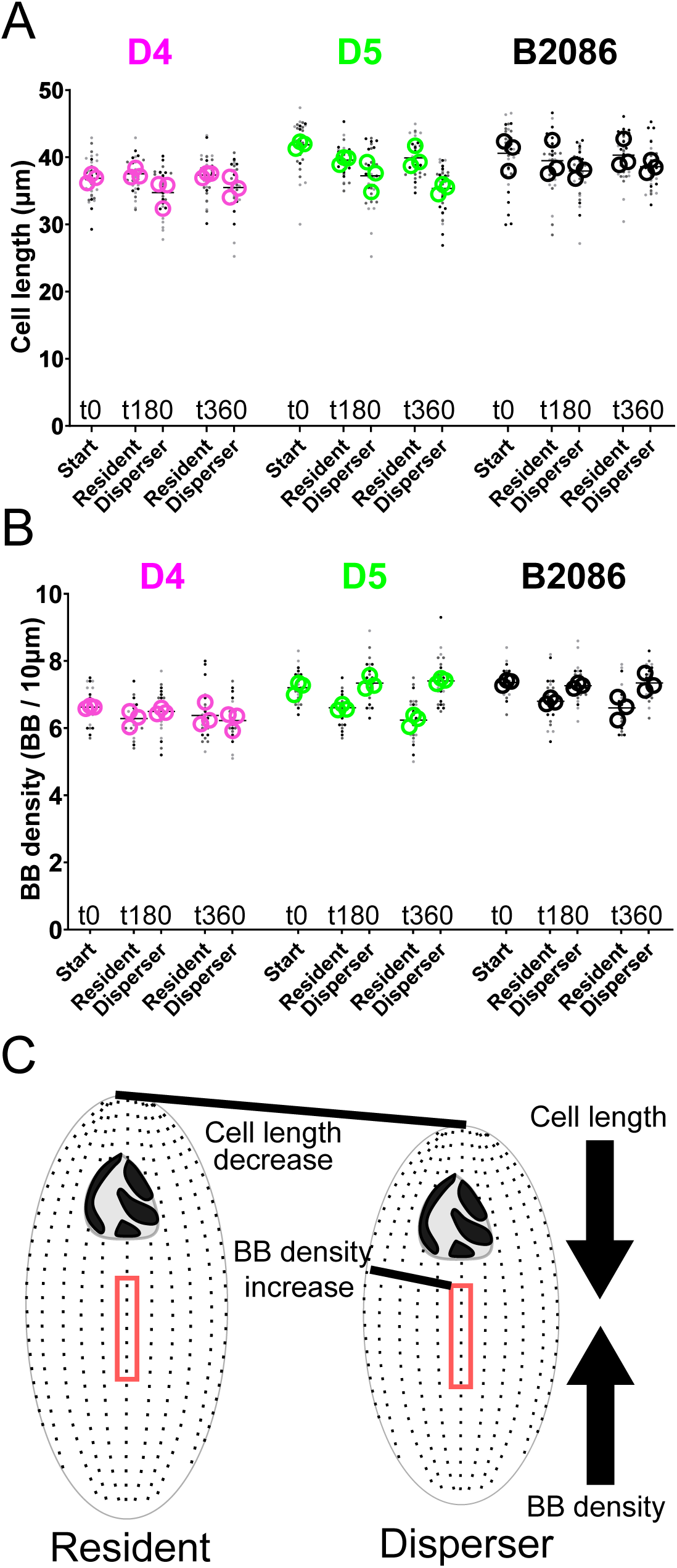
BB and cilia density and cell length increase in higher dispersing cells. (A) Cell length in D4 disperser cells relative to resident cells is reduced at 180 (p<0.0001) and 360 (p=0.02) min, cell length decreases in D5 disperser cells at 180 and 360 (p<0.0001) min. Cell length remains the same in B2086 cells at 180 (p=0.09) min and decreases at 360 (p=0.03) min. Data is represented as strain mean (black bar), the experimental means (hollow circles), and individual cell length (dots). (B) BB density in D4 disperser cells relative to resident cells is unchanged at 180 min (p=0.15) and 360 min (p=0.33). BB density is increased in D5 disperser cells at 180 and 360 min (p<0.0001). BB density is increased in B2086 disperser cells at 180 and 360 min (p<0.0001). Data is represented as strain mean (black bar), the experimental means (hollow circles), and mean cell BB density (dots). (C) Schematic for how a decrease in cell length can increase BB (and ciliary) density.

### Starvation promotes plastic changes to cell swimming and shape that trigger dispersal

Starvation is predicted to enhance dispersal in some strains, and triggers *T. thermophila* cells to become narrower and swim 2-3-fold faster ^24, 39^. We next tested whether dispersal promoting cell changes are induced by the environmental signal of starvation, and whether this response depends on the innate dispersal propensity of each strain. To test for the effect of starvation on plastic dispersal changes according to innate dispersal propensity, cells from D4 (low dispersal), D5 (high dispersal), and B2086 (very high dispersal) were starved by replacing growth media (1X SPP) with 10 mM Tris-HCl to induce “extreme disperser” phenotypes ^39, 40^.

A peak of fast cell swimming was observed for D5 and B2086 240 min after starvation (Figure S4A). Analyses were thus conducted to compare fed cultures with those that were starved for 240 min (Figure 4A). The rate of swimming increased for each strain (Figure 4B). The maximum swim speed of D5 (944 µm/sec) and B2086 (941 µm/sec) increased more than D4 (332 µm/sec). Moreover, the variation in swim speed between cells was greater in D5 and B2086 compared to D4. The standard deviation and coefficient of variation (%CV) for D4, D5 and B2086 was 60 µm/sec (25 %CV), 199 µm/sec (87 %CV) and 154 µm/sec (49 %CV), respectively. In summary, starvation caused all strains to increase their swimming speed, but D5 and B2086 exhibited a greater maximum change (2.8-fold) and cell-to-cell variation in swim speed compared to D4 cells.

**Figure 4.**
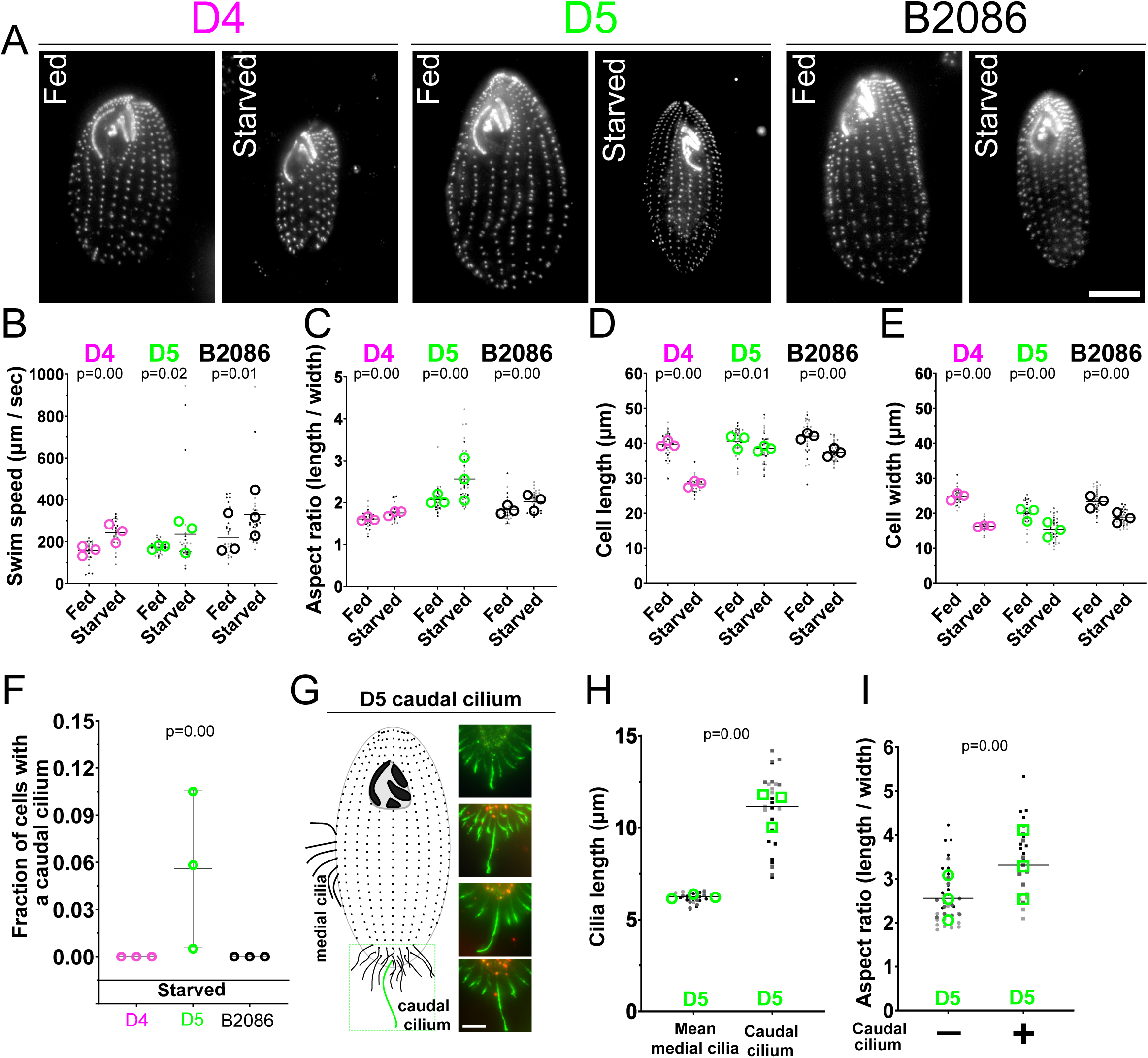
Starvation-induced dispersal increases BB density and formation of a caudal cilium. (A) BB localization (α-TtCen1; greyscale) and cell morphology in D4, D5 and B2086 cells from cycling (fed) and starved conditions (scale bar, 10µm). (B) Starved D4, D5, and B2086 cells swim faster than fed cells (p<0.0001, 0.02, and 0.01, respectively). Maximum speeds of both B2086 and D5 are 2.8-fold faster than the maximum speed of D4. Data is represented as strain mean (black bar), the experimental means (hollow circles), and individual cell speed (dots). (C) Starved cells increase in cell aspect ratio after starvation for all strains (p<0.0001 for D4, D5 and B2086). Data is represented as strain mean (black bar), the experimental means (hollow circles), and individual cell aspect ratio (dots). (D) Cell length decreases in all strains after starvation (D4, p<0.0001; D5, p=0.01; B2086, p<0.0001). Data is represented as strain mean (black bar), the experimental means (hollow circles), and individual cell length (dots). (E) Cell width decreases after starvation (p<0.0001 for D4, D5 and B2086). Data is represented as strain mean (black bar), the experimental means (hollow circles), and individual cell width (dots). (F) Starved D4 and B2086 cells do not generate a caudal cilium while starved D5 cells exhibit a subpopulation of cells with a caudal cilium (p<0.0001). Data is represented as strain mean and standard deviation (black bars), and the experimental means (hollow circles). (G) Model of D5 cell and cilia location (left). Fluorescence images of the caudal or cell posterior end in starved D5 cells. BBs are stained for BBs (α-TtCen1, red) and cilia (α -glutamylated and α-α-tubulin; green) (scale bar, 5 µm). (H) The caudal cilium is longer than the average cortical medial cilium (p<0.0001). Data is represented as a morphotype mean (black bar), the experimental means (hollow circles/squares), and mean cilia length of individual cells (dots). (I) Starved D5 cells with a caudal cilium have an increased aspect ratio (narrow cell shape) compared to starved D5 cells without a caudal cilium (p<0.0001). Data is represented as the morphotype mean (black bar), the experimental means (hollow circles/squares), and the aspect ratio of individual cells (dots).

We next analyzed the cell size and shape differences between D4, D5, and B2086 cells (Figure 4A). Narrow cell shape is associated with increased dispersal and motility ^4, 24, 26, 30, 47^. Consistently, in response to starvation, the three strains increased their aspect ratio (8%, 19% and 11%, respectively, for D4, D5 and B2086, Figure 4C). This occurred in D4 cells by a dramatic reduction of both cell length (29%) and width (34%), in D5 cells by a small reduction in cell length (5%) and a large reduction of cell width (23%), and in B2086 cells by a small reduction in cell length (11%) and a greater reduction in cell width (20%) (Figures 4D-E). Thus, general cell morphology changes in response to starvation are different between each strain.

### Increased BB density and caudal cilia are associated with dispersal propensity

To determine the relationship between cell swim speed and cilia, we quantified differences in BBs and cilia between the D4, D5, and B2086 cells after starvation for 240 min. No difference in the number of ciliary rows between fed and starved cells was observed. Combined with the decreases in cell width (D4=34%, D5=23%, B2086=20%), the lateral spacing between rows of BBs and cilia correspondingly decreased resulting in an increase in lateral BB density (Figure 4E and S4B). This is consistent with original starvation analyses^39^. The BB density within ciliary rows in each strain also uniquely responded to starvation (Figure S4C): D4 cells exhibited a 10% reduction in BB density, D5 cells had no change and B2086 cells increased BB density by 18%. This demonstrates that BB density in starvation increases as strain dispersal propensity increases.

No differences in cilia length were detected between fed and starved cells (Figure S4D). This suggests that cilia length is not responsive to starvation, exactly as cilia length does not change between residents and dispersers (see above). Interestingly, a small fraction of starved D5 cells, but not D4 or B2086 cells, possess a single caudal cilium (Figure 4F-G) that is 1.2-2.3-fold longer than the average medial cilium length (Figure 4H). The subpopulation of starved D5 cells with a caudal cilium are longer (16%, Figure S4E) and narrower (23%, Figure 4I and Figure S4FJ-K). These caudal cilia, only described once before in *T. thermophila*, do not beat and may serve as a rudder for steering motility (Movie 1; ^39^). The presence of a caudal cilium in a fraction of the starved D5 cells and their associated distinct morphology further supports the importance of plasticity in the dispersal process.

## Discussion

We have used an experimental approach linking ecological and cell biology competencies to determine how innate and plastic changes to fine-scale cellular activities can impact the dispersal process. Deciphering the mechanisms underlying a repertoire of intra-specific dispersal strategies is a critical step to understand dispersal evolution (e.g., how species modulate their dispersal response when facing environmental changes, or how variability in dispersal syndromes impacts ecosystem function or metapopulation dynamics)^42, 49^.

### Innate and plastic mechanisms underlie variation in dispersal propensity

*T. thermophila* have distinct intra-specific diversity in dispersal propensity and motility associated morphologies. Cell aspect ratio and cilia length predict the linearity of dispersers’ motility. Because no innate cell or cilia morphology directly correlates with the rate of dispersal, we suggest that different *T. thermophila* isolates implement different strategies to disperse. This diversity of strategies might result from tradeoffs between dispersal behaviors (e.g., linearity in motility) and efficiency in motility involving cell shape, cilia length and BB density, and/or colonization ability^21^. Interestingly, innate swim speed does not correlate with dispersal rate, nor with dispersers’ swimming characteristics. This absence of correlation is expected when dispersers’ movement dynamics result from plastic variability during dispersal, as was previously described in *T. thermophila*^26, 41^. Thus, we suggest that dispersal rates as measured in two-patch systems are not a simple by-product of random movements of each strain within a patch, but rather result at least in part from phenotypic characteristics specifically expressed during the dispersal process. Accordingly, during the dispersal and starvation experiments, cells develop new morphologies that are not present in the original population including in narrow cell shape, increased BB and cilia density, and a caudal cilium. Both cell shape and increased BB and cilia density are associated with increased phenotypic variance after the dispersal assays. For instance, when dispersal associated behavior is triggered by starvation, higher dispersing strains in standard conditions display fast but variable swim speeds, narrow cell shape, increased BB density, and/or a caudal cilium. Thus, our study highlights that phenotypic plasticity plays a major role in the morphological determinants of *T. thermophila* dispersal.

Dispersal is an inherently costly process involving movement through a matrix that separates habitats^50^. A morphology that improves the dispersal propensity of an individual may be detrimental to other fundamental processes like nutrient acquisition. To manage these tradeoffs, one strategy is to plastically activate a costly morphology in a context-dependent manner^15, 51^. Another strategy is to use existing morphological variation in a population where only a fraction of individuals exhibits potentially costly morphologies that are beneficial to dispersal. Both models can produce populations of individuals with unique dispersal capacity, consistent with the observations in this study. Moreover, this provides a morphological basis for dispersal propensity independent from a generalized innate morphology. We have shown in *T. thermophila* that plasticity mainly explains the variability in dispersal strategies observed (cells in resident, dispersing, or starved populations have morphologies that are not present in the original starting population). However, variation could also be generated through asymmetric cell division events that can produce unique cellular morphologies. In this study, we minimize this source of variation by restricting the dispersal time to four hours (less than one generation), six hours (less than two generations), or by pausing the cell cycle through starvation. While innate population variation may play a slight role in dispersal, our studies suggest that plastic changes of cell morphology that are transiently activated before or during dispersal is the main promotor of dispersal morphologies.

Starvation is generally a strong environmental cue that activates dispersal behavior, especially in microorganisms^52^. We observe that strains that proficiently disperse in fed conditions activate highly variable swimming behavior in response to starvation. Rather than an entire population synchronizing its motility rate when exposed to starvation, a spectrum of individuals with high motility is produced in strains with high dispersal propensity. The observed variation in swim speed could rely on multiple changes including gross cell morphology, BB and cilia spacing, ciliary dynamics, or metabolic output. We suspect that combinatorial effects, with differences in BB and ciliary morphology could explain the high variation in swim speeds measured for strains D5 and B2086. These strains increased their phenotypic heterogeneity in response to starvation stress through phenotypic plasticity. Such a diversifying phenotypic plasticity could be evolutionarily viewed as a risk spreading strategy in fluctuating and discontinuous environments allowing for highly heterogeneous dispersal behaviors in cells with the same genotype.

### Plasticity in cilia density and cell aspect ratio occurs during dispersal

Motility is an essential component of active dispersal. In most aquatic eukaryotic microorganisms, including *T. thermophila*, motility is driven by cilia-dependent fluid flow^53^. For multiple cilia to efficiently produce fluid flow, they are properly spaced relative to one another^18, 29, 30, 46, 47, 54^. In our dispersal assays, decreases in cell length and increases in BB and cilia density in highly dispersive strains were observed in cells that have successfully dispersed. This is consistent with *T. thermophila’s* BB counting mechanism which ensures the number of BBs in each cell remains nearly constant^18^. Here, a reduction in cell length will increase BB density. An increase in BB density is expected to increase cilia-dependent fluid flow and motility ^29, 30, 46, 47^. For comparison, *T. thermophila* genetic mutants that cause BB instability often display 15-35% decreases in BB density and exhibit reduced cell swimming ^34–36^. In this study, highly dispersive D5 and B2086 strains have increased BB density and we suggest this to be what improves their cilia-dependent cell swimming during dispersal. We suggest that the plastic increase in BB and cilia density in ciliated microorganisms that control BB frequency increases effective cell motility during dispersal.

The cell’s global geometry is also likely to impact cell motility^30, 47^. Elevated cell aspect ratio creates cells that are elongated relative to their width. The increased aspect ratio that is triggered by nutrient removal is consistent with strain dispersal rates and the general finding of associations between dispersal and narrow cell shape. However, passive plastic changes in cell volume could also occur because of changes to osmotic pressure during starvation. We suspect that combinatorial effects, with differences in BB and ciliary morphology could explain the high variation in swim speeds measured for the efficient dispersing strains D5 and B2086. It is plausible that narrow cell shape is an environment-dependent change that either directly impacts motility through effects on ciliary motility or reflects a metabolic state of the cell that promotes dispersal behaviors.

### The caudal cilium is a determinant of rapid cell motility

The caudal cilium is a transient structure that we observed in this study upon exposure to starvation. It is present only in a sub-population of starved D5 cells and is not found in fed cells. Moreover, the caudal cilium is associated with variable fast motility of cells. This is a distinct example of a transiently emerging morphology that is not present before dispersal behavior is initiated. Caudal cilia do not appear to beat and may serve as a rudder for steering rapid cell motility (Movie 1; ^39^). Caudal cilia have also been detected on other *Tetrahymena* species and ciliates including *Ichthyophthirius*, *Uronema*, *Paramecium* and *Coleps/Levicoleps* ^40, 55–60^. *Ichthyophthirius*, a parasitic ciliate closely related to *T. thermophila*, has distinct morphologies during its life cycle and possesses a caudal cilium only during its free-swimming dispersal stage prior to host invasion ^60^. The transient development of a caudal cilium further supports the importance of morphological plasticity and variation in dispersal. We posit that plasticity and variation in morphology is critical for managing the costs and risks surrounding the dispersal process.

## Conclusion

By decrypting the morphological determinants of dispersal at a previously unseen cellular level, we show that the dispersal process does not rely on innate phenotypic stereotypes. *T. thermophila* rather employs differential transient cell and ciliary morphologies to promote motility for dispersal, which could be important in managing energy cost trade-offs between morphologies that promote feeding and the transition to morphologies employed for efficient dispersal ^30, 47, 50, 61^. To determine if this link between cilia-dependent motility and dispersal is ubiquitous, future studies should now be developed in other taxa inhabiting ponds and lakes, but also river, ocean, and terrestrial biotas. Future studies should also elaborate on the molecular determinants, costs, and benefits of this panel of dispersal strategies, as well as their ecological and evolutionary consequences in real landscapes both when individuals colonize new patch (as in our study) and settle in an already occupied patch.

### Limitations of the study

We studied dispersal within highly simplified microcosms. The limitations of artificial systems, especially the generalization of the corresponding observations to nature, is a long-standing matter of concern in scientific research, but their obvious advantages lead to their massive use ^62, 63^. In the context of dispersal, behaviors appear to be similar in nature and in artificial systems for some species^63–66^. *In Tetrahymena*, prior studies from our groups and others have shown that dispersal behavior studied in the simple two-patch systems are congruent with what is known for other species in natura. For instance, cells adjust dispersal decisions relative to kinship, population density, resources, and temperature within these simplified microcosms^20–23, 42, 43^. However, techniques are not available to study the *T. thermophila* dispersal behaviors in natura, as for most microorganisms. Yet, our system is extremely simplified (axenic media, only two patches, absence of sex, only clonemates with a likely unreasonable cell density) and therefore far from natural conditions. The severity of some of these limitations could be easily evaluated, e.g., feeding *T. thermophila* with a single or a diversity of bacteria, introducing top predators and/or competing species, working on a heterogeneous populations of *T. thermophila*, or having multiple patches of different sizes and in physical different conditions. This would test whether dispersal and its related morphologies are dependent on specific environmental conditions. Nonetheless, in the long run, studies *in natura* will be necessary

The morphological analysis of *Tetrahymena* cells, BBs and cilia requires cell fixation and preparation for light microscopy. This can impact natural morphologies and subtle changes to BB and cilia morphology are to be expected. Moreover, this study measured a limited number of morphological parameters that can be expanded upon to further explore the diversity in changes with dispersal. These include, but are not limited to, cortical rigidity, oral apparatus positioning, ciliary beat frequency, waveform and metachronicity, mechanical coupling between BBs, and number and volume of vacuoles. In addition, our studies highlight a subset of the population and examples like the caudal cilium suggest that we may not accurately estimate the variance of morphological diversity in these systems. The low frequency and high variation of caudal cilia appearance in starved D5 cells make it challenging to study the functional role of this transient cilium. To image live cells with a caudal cilium, cells were slowed using viscous media and imaged when cells swam just above the cover glass. While the caudal cilium appears to be immotile compared to neighboring cilia, subtle beating may be difficult to visualize. Future quantitative studies will use automated approaches analyzing cell morphologies, and cilia dynamics to describe morphological and behavioral variance.

## ACKNOWLEDGEMENTS

AJ and CGP were supported by NIH/NHLBI # F31HL1474495 and NIH/NIGMS # R01GM099820. SJ, HP, and DL are part of the French Laboratory of Excellence project TULIP (ANR-10-LABX-41). CGP was supported by the 2018 Visiting Scientist program of French Laboratory of Excellence project TULIP (ANR-10-LABX-41). We thank Alex Stemm-Wolf for critical comments on the manuscript. SJ, HP and DL warmly thank Michèle Huet for the breeding of strains and her precious experimental advice.

## AUTHOR CONTRIBUTIONS

AJ, SJ, DL, and CGP designed the study and analyzed experimental data. AJ, SJ, and DL performed the experiments and collected the data. AJ and SJ conducted the computational analyses. HP provided conceptual and experimental assistance. All authors contributed substantially to the writing and revisions of this manuscript.

## Supplemental Movies

Movie 1: D5 cell with a caudal cilium.

Caudal cilium at the posterior end of D5 cell does not exhibit ciliary beating as observed in neighboring cortical cilia.

## Materials and Methods

### Tetrahymena cell culture

*Tetrahymena thermophila* D1-D22 strains were kindly provided by Paul Doerder (Cleveland State University, see Figure S1H) and B2086 was obtained from the *Tetrahymena* Stock Center (tetrahymena.vet.cornell.edu/index). Cultures for D1-D22 were maintained in 24-well culture plates by passaging approximately every 10 days, with two periods of cryopreservation. Each strain was grown independently and cultured in 0.6% SPP media (0.6% protease peptone, 0.03% yeast extract, 0.06% glucose, and 0.0009% Fe-EDTA) at 23°C. Cells collected for analysis were grown to mid-log phase (approximately 3×10^5^ cells/mL). Cell counts were determined using a Coulter Counter Z1 (Beckman Coulter), except for the quantification of dispersal rates (see below).

### Quantitative analyses of *Tetrahymena* cell swimming and morphology

Swim speed was calculated from 5 sec videos acquired at 20 frames per sec, individual cell swim paths were followed (for 4-10 frames) to determine average speed in µm/second (Figure S1B). The swim speed of cells is defined as the total distance travelled by cells divided by the duration of the trajectory, and linearity is the ratio between the net distance travelled (Euclidian distance between start and end positions), and the total distance travelled, such that higher values indicate straighter trajectories (Figure S1B). Cell length was measured as the distance from the anterior tip to the posterior end of the cell (Figure S1C). Cell width was measured as the distance across the cell’s dorsal-ventral axis perpendicular to the anterior-posterior cell axis, 1 μm posterior to the cell’s oral apparatus (Figure S1D). Aspect ratio was calculated as the length/width of the cell (Figure S1E). BB density was measured as the number of BBs (α-TtCen1) in a 10 µm distance of a ciliary row within the medial region of the cell (middle 50% of anterior-posterior axis) (Figure S1F). Ciliary rows per cell were counted through full cell z-stacks as the total number of individual rows of BBs around the cell’s lateral circumference (Figure S1G). Cilia length was measured using a segmented line as the distance from the distal end of the BB (α-TtCen1) to the tip of the cilium (α-–α-tubulin and α-glutamylated tubulin) (Figure S1H). Each of these measure parameters were used to generate an average morphology for all strains analyzed (Figure S1I).

### Dispersal

We performed dispersal assays to characterize the 22 *T. thermophila* strains using two-patch dispersal systems consisting of 1.5 mL tubes connected by a corridor made of a 2.5 cm long silicone tube with a 4 mm internal diameter ^20, 21, 67^. In *T. thermophila*, cells adjust dispersal decisions relative to kinship, population density, resources, and temperature within these simplified microcosms^20–23, 42, 43^. Such results can be observed because two-patch systems accurately mimic simplified metapopulations, i.e., suitable patches separated by a corridor (over 1000 times the size of a *T. thermophila* cell), with each patch being of sufficient size to host very large population sizes (up to ∼500 000 cells per patch; ^42^). Importantly, dispersal movements in these experimental microcosms influence *T. thermophila* local adaptation by generating gene flow among patches and affecting population differentiation^22^. The movements we quantify in these microcosms therefore match the classical definition of dispersal movements: movements between populations than can lead to gene flow^1, 2^. The dispersal system was filled with 4 mL of 0.6% SPP media and the central corridor was clamped. 100,000 cells were added to the resident patch. After 30 minutes of acclimatization, the corridor clamp was removed, and cells were allowed to disperse to the disperser patch for 4 hours at 23°C. After the 4 hrs, the corridor was clamped again and cells from each patch were collected, counted, and/or analyzed for swimming parameters described above. Dispersal rate was measured as the fraction of dispersers / total number of cells, as measured from 2 x 10 μL pipetted from each tube into chambers of a multichambered counting slide (Kima precision cell 301890). We immediately acquired 20 sec live cell movies from each chamber under dark-field microscopy to count cells and quantify movement characteristics using BEMOVI R-package ^68^.

To quantify swimming and morphology changes associated with dispersal in D4, D5 and B2086, large two-patch systems with more cells were required to perform immunofluorescence experiments. Dispersal assays were conducted using a two-patch system consisting of 5 mL habitat microtubes connected by an 8 cm long silicone tube corridor with a 4 mm internal diameter. This dispersal system was optimized for the increase in habitat size and corridor length (Figure S3A-D). This dispersal system was filled with 11 mL of 0.6% SPP media and the central corridor was clamped. 400,000 cells were added to the resident patch. As above, after acclimatization, the corridor clamp was removed, and cells were allowed to disperse to the disperser patch for 6 hours at 30°C.

### Starvation

Starvation experiments were conducted by growing cells in 5 mL cultures with 2% SPP growth media at 30°C to mid log (approximately 3×10^5^ cells/mL). Cells were centrifuged in 10 mM Tris HCl pH 7.4, then washed in 10 mM Tris HCl pH 7.4 again before resuspending cells (3×10^5^ cells/mL) in 5 mL of 10 mM Tris HCl pH 7.4. Cells were incubated 240 min at 30°C before collection and analysis. Analysis of swim speed was conducted as described above, standard deviation and percent coefficient of variation (% CV) was calculated in Graphpad Prism on the mean speeds of individual cells.

Visualization of the caudal cilium in live D5 cells was performed by pelleting cells at 0.6 xg in a microfuge and resuspending them in 2% polyethylene oxide (PEO, molecular weight 900,000, Acros Organics) dissolved in 10 mM Tris pH 7.4^27^. The PEO solution increases viscosity and slows mobility for visualization. DIC movies were acquired at 100 frames per sec (10 ms exposure).

### Sample preparation for immunofluorescence

Immunofluorescence imaging procedures were modified from prior studies to maximize the quality of BB, cilia staining and retention of cell shape^37^. For BB and cilia visualization, cells were washed in 10 mM Tris HCl pH 7.4, followed by PHEM buffer (60 mM 1,4-piperazinediethanesulfonic acid, 25 mM 4-(2-hydroxyethyl)-1-piperazineethanesulfonic acid, 10 mM EGTA, and 2 mM MgCl_2_, pH 6.9), and then fixed in a Triton X-100 / paraformaldehyde solution (0.25% Triton X-100 and 3.2% paraformaldehyde in PHEM buffer) for 5 min. Cells were then incubated in Triton X-100 solution (0.25% Triton X-100 in PHEM buffer) for 10 min on ice. They were washed three times in 0.5% bovine serum albumin (BSA) in PBS (BSA-PBS) before a 24 hr incubation in primary antibody diluted in 0.5% BSA-PBS at 4°C. They were then washed three times in 0.5% BSA-PBS before incubation in secondary antibody diluted in 0.5 % BSA-PBS at 23°C for 2 hours. They were again washed three times in 0.5% BSA-PBS and pelleted. 2 µL of cells from the pellet were added to a coverslip and mounted in 6.5 µL Citifluor mounting media (Ted Pella). Samples were then sealed using clear nail polish.

The primary antibodies used in this study were α-TtCen1 (1:1000 dilution ^69^), α-glutamylation (1:1000 dilution, GT335, Adipogen AG-20B-0020-C100 ^54^), and α-α-tubulin (1:200 dilution, 12G10, DSHB AB_1157911 ^70^). Secondary antibodies (Alexa Fluor 488 (Invitrogen A11034), Alexa Fluor 594 (Invitrogen A32740), or Alexa Fluor 647 (Invitrogen A21244) goat α–rabbit IgG, and Alexa Fluor 488 (Invitrogen A11029), Alexa Fluor 594 (Invitrogen A32742), or Alexa Fluor 647 (Invitrogen A21236) goat α–mouse IgG) were used at 1:1000 dilution.

### Microscopy

Fluorescence imaging was performed as previously described ^37^. Briefly, either a Nikon wide-field TiE fluorescence microscope stand equipped with a 100x NA 1.4 Plan Apo objective and an Andor Xyla 4.2 CMOS camera or a Nikon X1 confocal TiE fluorescence microscope stand equipped with a 100x NA 1.4 Plan Apo objective and an Andor iXon EMCCD camera were used depending upon the experiment. Images were acquired using NIS Elements software. All fluorescence imaging was conducted at approximately 25°C and exposure times were between 50-500 msec, depending on the experiment. Only *T. thermophila* cells in Stage 1 (G1) of cell cycle and cells that were not visibly damaged by the fixation and staining process were included in the analyses.

### Statistics and computational analysis

All experimental data sets represent a minimum of three biological replicates. For morphological analysis, each replicate has approximately 10 cells (at least 30 cells per experimental condition) unless otherwise indicated in the Figure Legends. Statistical *p*-values are included in each data set in the corresponding Figure Legend. Statistical tests were run in Prism8 (GraphPad Software). Normally distributed continuous data sets were analyzed using unpaired, two-tailed Student’s t-test. Non-normally distributed continuous data sets were analyzed using unpaired, two-tailed Mann-Whitney test. Data sets with multiple comparisons were analyzed using one-way ANOVA. All *p*-values are numerically presented to two decimal places. Lines and bars on all plots indicate the mean and standard deviation, respectively. For correlations between traits, each measured parameter was ranked among the isolates and simple linear regression was used to compare the parameters (Figure 2B). Parameters that correlated had a *F*-test p-value for a non-zero slope of < 0.05 and were assigned a value of 1. Parameters that did not correlate had p-value for a non-zero slope of > 0.05 and were assigned a value of 0. Standard deviation of residents and dispersers from the start populations was calculated as the standard deviation of each strain’s resident or disperser measurements to their start population’s mean (Figure S3E,F). One-way ANOVA for each strain with Dunett’s multiple comparisons to the start condition mean was conducted and reported in the figure legend. The fraction of start overlap to simulation means was calculated using the fraction of start condition measurements that fell within the range of their corresponding resident or disperser 95% confidence intervals (Figure S3 G, H). This is used as an approximation of morphological contribution of innate start population morphology to the mean morphology observed in corresponding residents and dispersers.

### Data Representation

Each data point (representing a single cell) is indicated as a single dot on the figure plots. Replicates for morphological data are denoted by different shades (light grey, dark grey and black), the mean of each replicate is denoted by hollow colored circles. Lines denote the mean for the entire experimental condition. All lines indicate the mean and all error bars indicate standard deviation.

---

**Figure S1.**
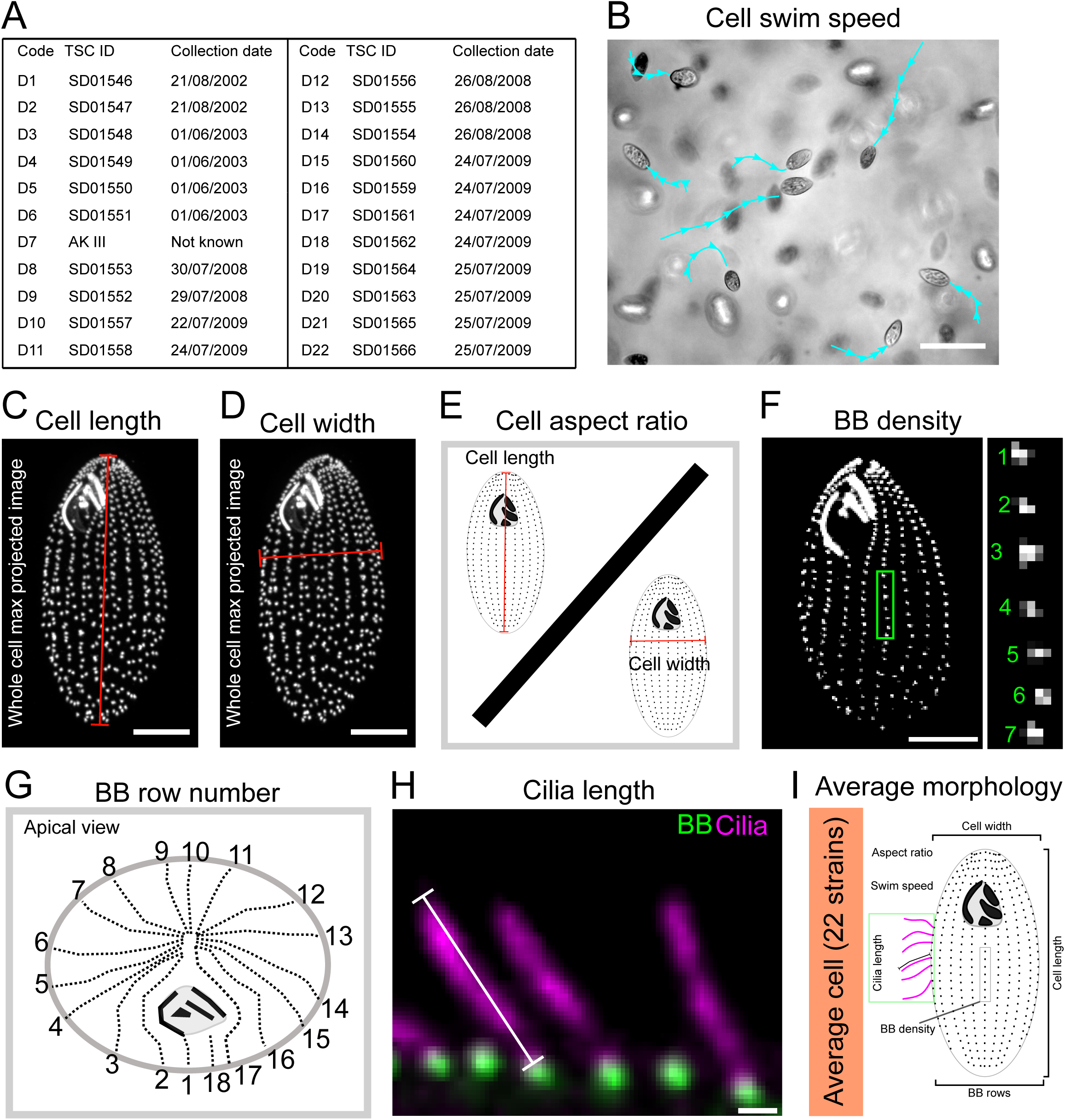
Measurement methods and parameters for *T. thermophila* strains. (A) We present here the codes of our 22 strains with the associated reference number of the Tetrahymena Stock Center (https://tetrahymena.vet.cornell.edu) and the dates at which Dr. Paul Doerder collected them in the field. (B-H) Graphical representations of the techniques used in the 22 *T. thermophila* strains to measure morphological parameters (see methods). (B scale bar, 100 µm, C, D, and F scale bars, 10 µm. H scale bar, 1 µm). (I) Graphical representations of all morphological parameters measured in each of the 22 *T. thermophila* strains.

**Figure S2.**
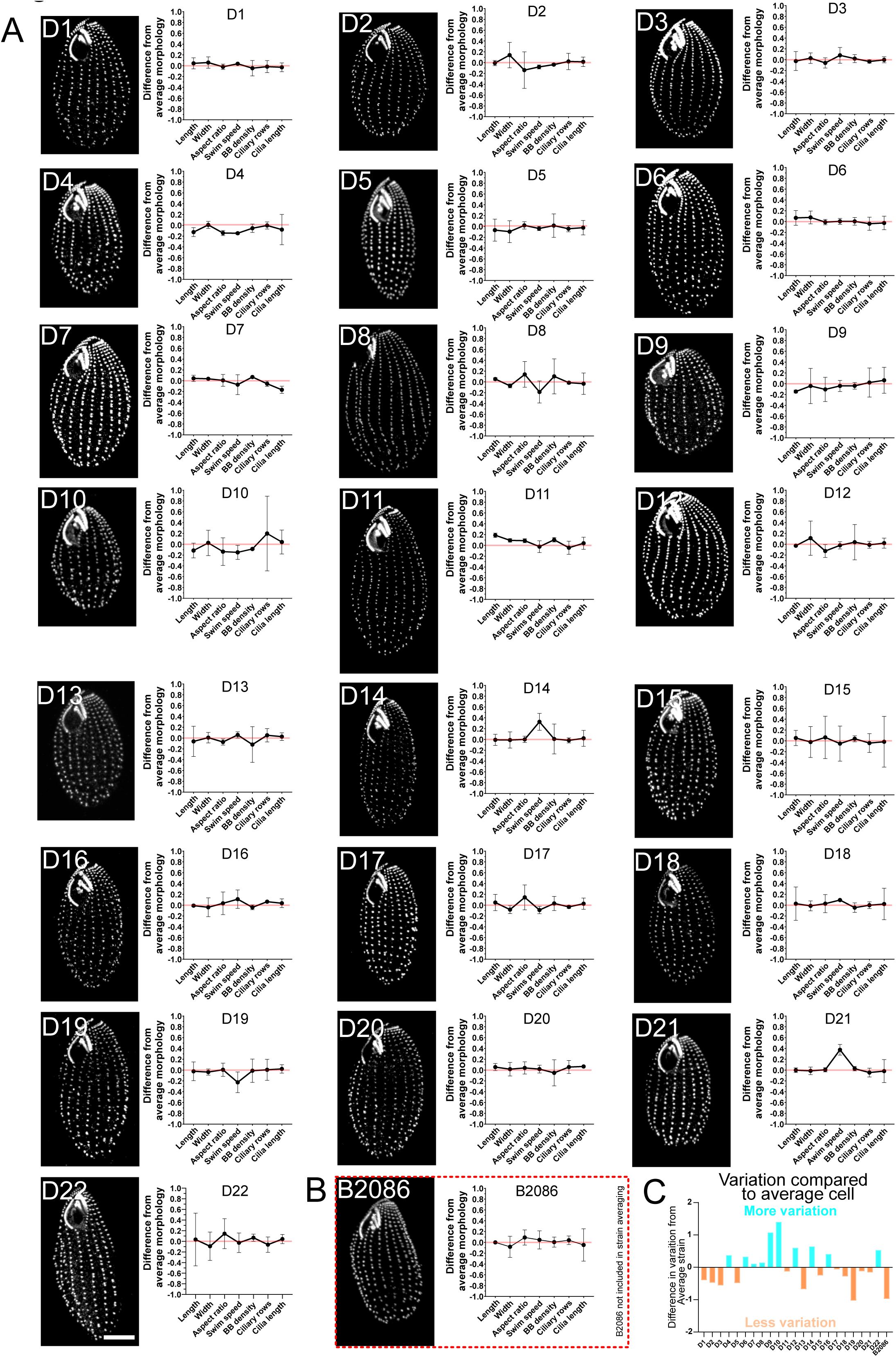
*T. thermophila* strains exhibit unique morphologies and levels of variation. (A) Representative BB (α-TtCen1; greyscale) and cell morphology for each of the 22 *T. thermophila* strains display unique morphological differences (scale bar, 10 µm). Graphs to the right of each image show the individual strain’s morphologies normalized to the inter-strain average (red line). (B) Representative BB (α-TtCen1; greyscale) and cell morphology of the B2086 lab strain B2086 (scale bar, 10 µm). Graph to the right of the image shows B2086 morphologies normalized to the inter-strain average (red dotted line). B2086 was not included in the inter-strain average. (C) Bar chart indicates the sum of standard deviations for all morphological parameters for individual strains normalized to the inter-strain mean. Red bars indicate the strains with values above the average morphological variation, blue bars indicate strains with values below the average morphological variation.

**Figure S3.**
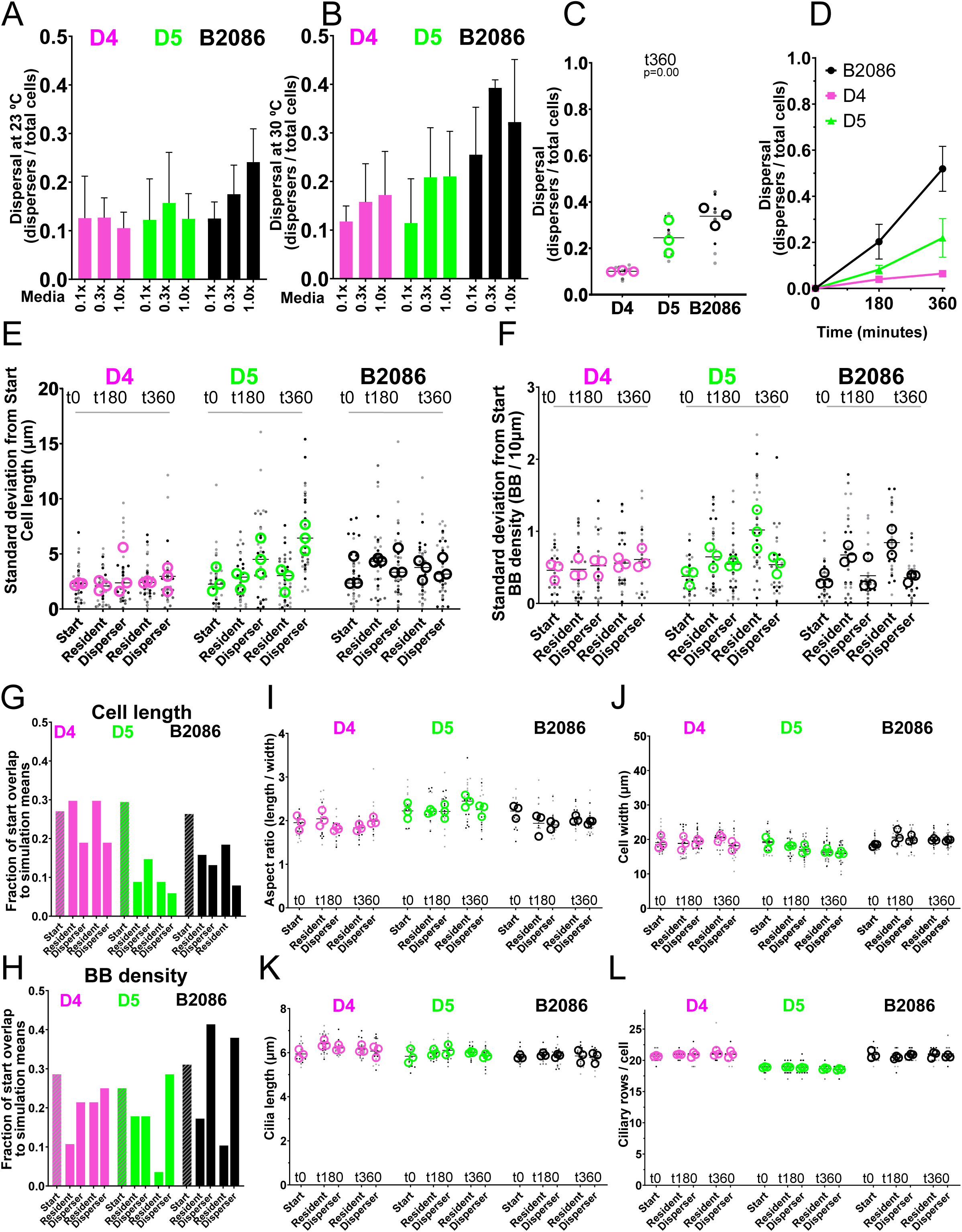
BB and cilia density and cell length increase in faster dispersing cells. (A) Dispersal rate over 6 hrs in two-patch system for low dispersal (D4), high dispersal (D5), and very high dispersal (B2086) at 23°C were conducted in SPP (growth media) concentrations of 0.1X, 0.3X and 1.0X. A consistent increase in dispersal in particular media concentration for all strains was not observed. Bars indicate means and error bars indicate standard deviation. (B) Dispersal rate over 6 hrs in two-patch system for low dispersal (D4), high dispersal (D5), and very high dispersal (B2086) at 30°C were conducted in SPP (growth media) concentrations of 0.1X, 0.3X and 1.0X. All three strain show slight preference for 0.3X or 1.0X media concentrations. Because higher dispersal fractions were observed at 30°C (compared to 23°C), all further dispersal experiments were conducted at 30°C in 0.3X SPP. Bars indicate means and error bars indicate standard deviation. (C) Dispersal rate over 6 hrs in two-patch system for low dispersal (D4), high dispersal (D5), and very high dispersal (B2086) *T. thermophila* strains show differences in dispersal (p<0.0001). Data is represented as the strain mean (black bar), the experimental means (hollow circles), and the technical replicates within experiments (dots). (D) Time course of dispersal rate at 0, 180 and 360 min in two-patch system for low dispersal (D4), high dispersal (D5), and very high dispersal (B2086) *T. thermophila* strains. Data is represented as the strain mean and standard deviation (colored circle and error bars) for 3 biological replicates. (E) Standard deviation in cell length between Start and resident and disperser cells is unchanged in D4 (ANOVA p=0.30, Dunett’s multiple comparison test p=0.98, 0.36, 0.99, 0.66). D5 t360 disperser cell length SD is higher (ANOVA p=<0.01, Dunett’s multiple comparison test p=0.99, <0.01, 0.99, <0.01). B2086 cell length SD is unchanged (ANOVA p=<0.33, Dunett’s multiple comparison test p=0.18, 0.33, 0.88, 0.89). Data is represented as the strain mean (black bar), the experimental means (hollow circles), and SD in cell length of individual cells (dots). (F) Standard deviation in BB density between Start and resident and disperser cells is unchanged in D4 (ANOVA p=0.51, Dunett’s multiple comparison test p=0.99, 0.96, 0.69, 0.33). D5 t360 resident BB density SD is higher (ANOVA p=<0.01, Dunett’s multiple comparison test p=0.06, 0.36, <0.01, 0.41). B2086 t180 and t360 resident BB density SD is lower (ANOVA p=<0.01, Dunett’s multiple comparison test p=<0.01, 0.93, <0.01, 0.99). Data is represented as the strain mean (black bar), the experimental means (hollow circles), and SD in BB density of individual cells (dots). (G) The fraction of all cell length measurements in the Start condition that overlap with condition means (defined as 95% confidence interval) in Start, t180 Resident, t180 Disperser, t360 Resident, t360 are represented as a bar graph. (H) The fraction of all cell BB density measurements in the Start condition that overlap with condition means (defined as 95% confidence interval) in Start, t180 Resident, t180 Disperser, t360 Resident, t360 are represented as a bar graph. (I) Cell aspect ratio in D4 disperser cells relative to residents is increased at 180 min (p<0.0001) and at 360 min (p=0.01), is unchanged in D5 disperser cells at 180 min (p=0.95) and increased at 360 min (p=0.02) and remains the same in B2086 cells at 180 min (p=0.65) and at 360 min (p=0.20). Data is represented as the strain mean (black bar), the experimental means (hollow circles), and the aspect ratio of individual cells (dots). (J) Cell width in D4 disperser cells relative to residents is unchanged at 180 min (p=0.18) and increased at 360 min (p<0.0001), cell width in D5 disperser cells is unchanged at 180 min (p=0.06) and 360 min (p=0.33) and remains the same in B2086 cells at 180 min (p=0.34) and 360 min (p=0.72). Data is represented as the strain mean (black bar), the experimental means (hollow circles), and the width of individual cells (dots). (K) Cilia length in D4 disperser cells relative to residents is increased at 180 min (p=0.03) and unchanged at 360 min (p=0.34), is unchanged in D5 disperser cells at 180 min (p=0.17) and is decreased at 360 min (p=0.04) and remains the same in B2086 cells at 180 min (p=0.57) and at 360 min (p=0.45). Data is represented as the strain mean (black bar), the experimental means (hollow circles), and the mean cilia length of individual cells (dots). (L) The number of ciliary rows in D4 disperser cells relative to residents is unchanged at 180 (p=0.63) and 360 (p=0.35) min, is unchanged in D5 disperser cells at 180 (p=0.54) and 360 (p=0.43) min and remains the same in B2086 cells at 180 min (p=0.07) and 360 (p=0.35) min. Data is represented as the strain mean (black bar), the experimental means (hollow circles), and the number of ciliary rows of individual cells (dots).

**Figure S4.**
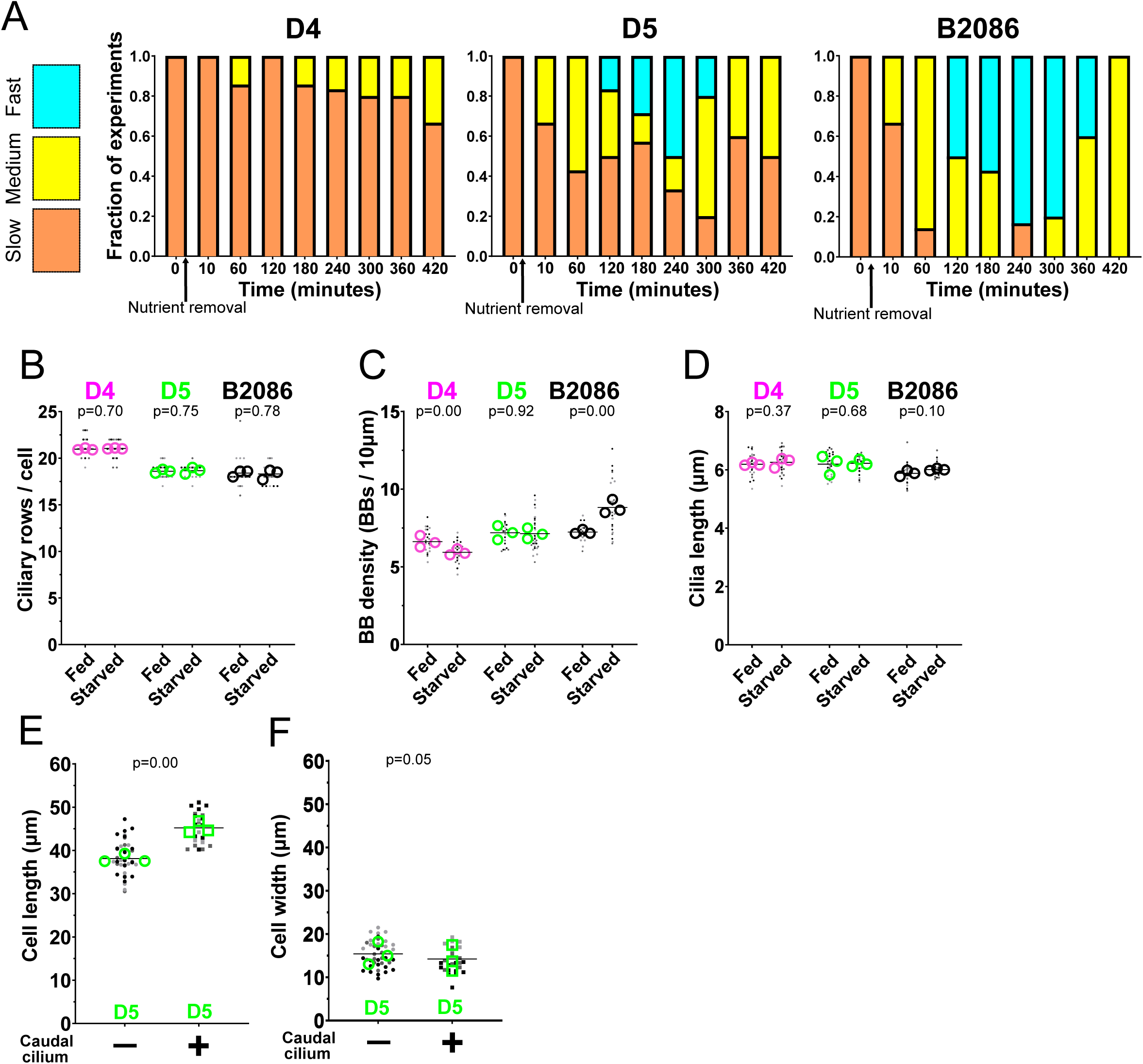
*T. thermophila* strains exhibit swimming responses to nutrient deprivation. (A) Starved cells were observed to determine when they first displayed fast swim speeds. For each timepoint, the relative swim speed for the entire population was visually judged and recorded to be either slow, medium, or fast. The most common timepoint where fast swimming was observed was 240 min post starvation (D4=0% of experiments, D5=50% of experiments, B2086 83% of experiments). Data is represented by colored bars that indicate the fraction of experiments (7 biological replicates) where cells with Slow (blue), Medium (green), or Fast (orange) swimming behaviors are observed. (B) Ciliary row number per cell does not change after starvation (D4, p=0.70; D5, p=0.75; B2086, p=0.78). Data is represented as strain mean (black bar), the experimental means (hollow circles), and the number of ciliary rows in an individual cell (dots). (C) BB density is differentially changed in each strain, D4 decreases (p<0.0001), D5 is unchanged (p=0.92), and B2086 increases (p<0.0001). Data is represented as strain mean (black bar), the experimental means (hollow circles), and mean BB density of an individual cell (dots). (D) Cilia length does not change after starvation for any strain (D4 p=0.37; D5 p=0.61; B2086 p=0.10). Data is represented as strain mean (black bar), the experimental means (hollow circles), and mean cilia length of an individual cell (dots). (E) Starved D5 cells possessing a caudal cilium are longer than starved D5 cells without a caudal cilium (p<0.0001). Data is represented as the morphotype mean (black bar), the experimental means (hollow circles/squares), and the length of individual cells (dots). (F) Starved D5 cells with a caudal cilium are 10% narrower than starved D5 cells without a caudal cilium (p=0.05). Data is represented as the morphotype mean (black bar), the experimental means (hollow circles/squares), and the width of individual cells (dots).

## REFERENCES

1. Clobert, J., et al., Dispersal Ecology and Evolution. 2012: OUP Oxford.

2. Ronce, O., How Does It Feel to Be Like a Rolling Stone? Ten Questions About Dispersal Evolution. Annual Review of Ecology, Evolution, and Systematics, 2007. 38(1): p. 231–253.

3. Van Dyck, H. and M. Baguette, Dispersal behaviour in fragmented landscapes: routine or special movements? Basic and Applied Ecology, 2005. 6(6): p. 535–545.

4. Stevens, V.M., et al., Dispersal syndromes and the use of life-histories to predict dispersal. Evol Appl, 2013. 6(4): p. 630–42.

5. Clobert, J., et al., Informed dispersal, heterogeneity in animal dispersal syndromes and the dynamics of spatially structured populations. Ecol Lett, 2009. 12(3): p. 197–209.

6. Snell, R.S., et al., Consequences of intraspecific variation in seed dispersal for plant demography, communities, evolution and global change. AoB PLANTS, 2019. 11(4).

7. Comte, L. and J.D. Olden, Evidence for dispersal syndromes in freshwater fishes. Proceedings of the Royal Society B: Biological Sciences, 2018. 285(1871): p. 20172214.

8. Le Galliard, J.F., et al., Patterns and processes of dispersal behaviour in arvicoline rodents. Molecular Ecology, 2012. 21(3): p. 505–523.

9. Bonte, D. and M. Saastamoinen, Dispersal syndromes in spiders and butterflies, in Dispersal ecology and evolution. 2012, Oxford University Press. p. 161–170.

10. Stevens, V.M., C. Turlure, and M. Baguette, A meta-analysis of dispersal in butterflies. Biological Reviews, 2010. 85(3): p. 625–642.

11. Cheptou, P.-O., et al., Rapid evolution of seed dispersal in an urban environment in the weed Crepis sancta. Proceedings of the National Academy of Sciences, 2008. 105(10): p. 3796–3799.

12. Ronce, O. and J. Clobert, Dispersal syndromes. Dispersal ecology and evolution, 2012. 155: p. 119–138.

13. Murren, C.J., et al., Constraints on the evolution of phenotypic plasticity: limits and costs of phenotype and plasticity. Heredity (Edinb), 2015. 115(4): p. 293–301.

14. Fox, R.J., et al., Beyond buying time: the role of plasticity in phenotypic adaptation to rapid environmental change. 2019, The Royal Society.

15. Cote, J., et al., Evolution of dispersal strategies and dispersal syndromes in fragmented landscapes. Ecography, 2017. 40(1): p. 56–73.

16. Bayless, B.A., D.F. Galati, and C.G. Pearson, Tetrahymena basal bodies. Cilia, 2015. 5: p. 1.

17. Nanney, D., S. Chen, and E. Meyer, Scalar constraints in Tetrahymena evolution. Quantitative basal body variations within and between species. The Journal of cell biology, 1978. 79(3): p. 727–736.

18. Nanney, D.L., Corticotype transmission in Tetrahymena. Genetics, 1966. 54(4): p. 955–68.

19. Doerder, F.P. and C. Brunk, Natural populations and inbred strains of Tetrahymena, in Methods in cell biology. 2012, Elsevier. p. 277–300.

20. Chaine, A.S., et al., Kin-based recognition and social aggregation in a ciliate. Evolution: International Journal of Organic Evolution, 2010. 64(5): p. 1290–1300.

21. Jacob, S., et al., Cooperation-mediated plasticity in dispersal and colonization. Evolution, 2016. 70(10): p. 2336–2345.

22. Jacob, S., et al., Gene flow favours local adaptation under habitat choice in ciliate microcosms. Nat Ecol Evol, 2017. 1(9): p. 1407–1410.

23. Jacob, S., et al., Habitat choice meets thermal specialization: Competition with specialists may drive suboptimal habitat preferences in generalists. Proc Natl Acad Sci U S A, 2018. 115(47): p. 11988–11993.

24. Fjerdingstad, E.J., et al., Evolution of dispersal and life history strategies--Tetrahymena ciliates. BMC Evol Biol, 2007. 7: p. 133.

25. Schtickzelle, N., et al., Cooperative social clusters are not destroyed by dispersal in a ciliate. BMC Evol Biol, 2009. 9: p. 251.

26. Pennekamp, F., J. Clobert, and N. Schtickzelle, The interplay between movement, morphology and dispersal in Tetrahymena ciliates. PeerJ, 2019. 7: p. e8197.

27. Galati, D.F., et al., DisAp-dependent striated fiber elongation is required to organize ciliary arrays. J Cell Biol, 2014. 207(6): p. 705–15.

28. Jiang, Y.Y., et al., LF4/MOK and a CDK-related kinase regulate the number and length of cilia in Tetrahymena. PLoS Genet, 2019. 15(7): p. e1008099.

29. Elgeti, J. and G. Gompper, Emergence of metachronal waves in cilia arrays. Proc Natl Acad Sci U S A, 2013. 110(12): p. 4470–5.

30. Omori, T., H. Ito, and T. Ishikawa, Swimming microorganisms acquire optimal efficiency with multiple cilia. Proc Natl Acad Sci U S A, 2020. 117(48): p. 30201–30207.

31. Bottier, M., et al., How Does Cilium Length Affect Beating? Biophys J, 2019. 116(7): p. 1292–1304.

32. Nanney, D.L. and J.W. McCoy, Characterization of the species of the Tetrahymena pyriformis complex. Trans Am Microsc Soc, 1976. 95(4): p. 664–82.

33. Wloga, D., et al., Members of the NIMA-related kinase family promote disassembly of cilia by multiple mechanisms. Mol Biol Cell, 2006. 17(6): p. 2799–810.

34. Pearson, C.G., et al., Basal body stability and ciliogenesis requires the conserved component Poc1. J Cell Biol, 2009. 187(6): p. 905–20.

35. Bayless, B.A., et al., Bld10/Cep135 stabilizes basal bodies to resist cilia-generated forces. Mol Biol Cell, 2012. 23(24): p. 4820–32.

36. Bayless, B.A., et al., Asymmetrically localized proteins stabilize basal bodies against ciliary beating forces. J Cell Biol, 2016. 215(4): p. 457–466.

37. Junker, A.D., et al., Microtubule glycylation promotes attachment of basal bodies to the cell cortex. J Cell Sci, 2019. 132(15).

38. Soh, A.W.J., et al., Ciliary force-responsive striated fibers promote basal body connections and cortical interactions. J Cell Biol, 2020. 219(1).

39. Nelsen, E.M., Transformation in *Tetrahymena thermophila*. Development of an inducible phenotype. Dev Biol, 1978. 66(1): p. 17–31.

40. Nelsen, E.M. and L.E. Debault, Transformation in Tetrahymena pyriformis: description of an inducible phenotype. J Protozool, 1978. 25(1): p. 113–9.

41. Jacob, S., et al., Fragmentation and the context-dependence of dispersal syndromes: matrix harshness modifies resident-disperser phenotypic differences in microcosms. Oikos, 2020. 129(2): p. 158–169.

42. Jacob, S., et al., Variability in Dispersal Syndromes Is a Key Driver of Metapopulation Dynamics in Experimental Microcosms. Am Nat, 2019. 194(5): p. 613–626.

43. Pennekamp, F., et al., Dispersal propensity in *Tetrahymena thermophila* ciliates - a reaction norm perspective. Evolution, 2014. 68(8): p. 2319–30.

44. Altermatt, F., et al., Big answers from small worlds: a user’s guide for protist microcosms as a model system in ecology and evolution. Methods in Ecology and Evolution, 2015. 6(2): p. 218–231.

45. Laurent, E., N. Schtickzelle, and S. Jacob, Fragmentation mediates thermal habitat choice in ciliate microcosms. Proc Biol Sci, 2020. 287(1919): p. 20192818.

46. Rode, S., J. Elgeti, and G. Gompper, Multi-ciliated microswimmers-metachronal coordination and helical swimming. Eur Phys J E Soft Matter, 2021. 44(6): p. 76.

47. Ito, H., T. Omori, and T. Ishikawa, Swimming mediated by ciliary beating: comparison with a squirmer model. Journal of Fluid Mechanics, 2019. 874: p. 774–796.

48. Galati, D.F., et al., Automated image analysis reveals the dynamic 3-dimensional organization of multi-ciliary arrays. Biol Open, 2015. 5(1): p. 20–31.

49. Little, C.J., E.A. Fronhofer, and F. Altermatt, Dispersal syndromes can impact ecosystem functioning in spatially structured freshwater populations. Biology letters, 2019. 15(3): p. 20180865.

50. Bonte, D., et al., Costs of dispersal. Biol Rev Camb Philos Soc, 2012. 87(2): p. 290–312.

51. Benard, M.F. and S.J. McCauley, Integrating across life-history stages: consequences of natal habitat effects on dispersal. The American Naturalist, 2008. 171(5): p. 553–567.

52. McDougald, D., et al., Should we stay or should we go: mechanisms and ecological consequences for biofilm dispersal. Nature Reviews Microbiology, 2012. 10(1): p. 39–50.

53. Mitchell, D.R., The evolution of eukaryotic cilia and flagella as motile and sensory organelles. Eukaryotic Membranes and Cytoskeleton, 2007: p. 130–140.

54. Wolff, A., et al., Distribution of glutamylated alpha and beta-tubulin in mouse tissues using a specific monoclonal antibody, GT335. Eur J Cell Biol, 1992. 59(2): p. 425–32.

55. Corliss, J., Tetrahymena paravorax n. sp., the first caudal-ciliated member of the genus referrable to the vorax-patula complex. J. Protozool., 1957. 4: p. 13.

56. Foissner, W., Y. Kusuoka, and S. Shimano, Morphology and gene sequence of Levicoleps biwae n. gen., n. sp. (Ciliophora, Prostomatida), a proposed endemic from the ancient Lake Biwa, Japan. J Eukaryot Microbiol, 2008. 55(3): p. 185–200.

57. Holz JR, G.G. and J.O. Corliss, Tetrahymena setifera n. sp., a member of the genus Tetrahymena with a caudal cilium. The Journal of Protozoology, 1956. 3(3): p. 112–118.

58. Lu, B.R., et al., Morphology and molecular phylogeny of two colepid species from China, Coleps amphacanthus Ehrenberg, 1833 and Levicoleps biwae jejuensis Chen et al., 2016 (Ciliophora, Prostomatida). Dongwuxue Yanjiu, 2016. 37(3): p. 176–85.

59. Tamm, S.L., Laser microbeam study of a rotary motor in termite flagellates. Evidence that the axostyle complex generates torque. J Cell Biol, 1978. 78(1): p. 76–92.

60. Kozel, T.R., Scanning electron microscopy of theronts of Ichthyophthirius multifiliis: their penetration into host tissues. Transactions of the American Microscopical Society, 1986: p. 357–364.

61. Gilpin, W., V.N. Prakash, and M. Prakash, Vortex arrays and ciliary tangles underlie the feeding–swimming trade-off in starfish larvae. Nature Physics, 2017. 13(4): p. 380–386.

62. Bestion, E., et al., Habitat fragmentation experiments on arthropods: what to do next? Current opinion in insect science, 2019. 35: p. 117–122.

63. Benton, T.G., et al., Microcosm experiments can inform global ecological problems. Trends Ecol Evol, 2007. 22(10): p. 516–21.

64. Legrand, D., et al., The Metatron: an experimental system to study dispersal and metaecosystems for terrestrial organisms. Nature methods, 2012. 9(8): p. 828–833.

65. Legrand, D., et al., Ranking the ecological causes of dispersal in a butterfly. Ecography, 2015. 38(8): p. 822–831.

66. Le Galliard, J.F., R. Ferrière, and J. Clobert, Effect of patch occupancy on immigration in the common lizard. Journal of Animal Ecology, 2005. 74(2): p. 241–249.

67. Jacob, S., et al., Social information from immigrants: multiple immigrant-based sources of information for dispersal decisions in a ciliate. J Anim Ecol, 2015. 84(5): p. 1373–83.

68. Pennekamp, F., N. Schtickzelle, and O.L. Petchey, BEMOVI, software for extracting behavior and morphology from videos, illustrated with analyses of microbes. Ecol Evol, 2015. 5(13): p. 2584–95.

69. Stemm-Wolf, A.J., et al., Basal body duplication and maintenance require one member of the *Tetrahymena thermophila* centrin gene family. Mol Biol Cell, 2005. 16(8): p. 3606–19.

70. Thazhath, R., C. Liu, and J. Gaertig, Polyglycylation domain of beta-tubulin maintains axonemal architecture and affects cytokinesis in Tetrahymena. Nat Cell Biol, 2002. 4(3): p. 256–9.

